# Half-match recombination drives bridge RNA-guided excision and off-target insertion

**DOI:** 10.64898/2026.08.31.748434

**Authors:** Bingliang Xie, Kuang Hu, Hengyi Yang, Jaymin R. Patel, Carlotta Ronda, Benjamin E. Rubin

**Author notes:** Corresponding authors (C.R.), (B.E.R.).

## Abstract

IS110-family bridge recombinases are a recently identified class of compact, RNA-guided editors in which a bridge RNA (bRNA) directs the recombination of a donor DNA into a target site. In the current model, the bRNA engages fully complementary donor and target sequences within a single synaptic complex to drive double-stranded recombination, implying that the transposon is cut from its donor site rather than copied, yet neither the strandedness of the excised intermediate nor the requirement for full complementarity has been tested directly. Here we reconstituted IS621 recombination in a cell-free transcription–translation system, building representative arrangements of the excision and insertion reactions and characterizing the outcomes. We find that IS621 predominantly excises a single strand, releasing a single-stranded circle and leaving the donor site intact, consistent with copy-and-paste transposition. By introducing mismatches into the bRNA target sequences, we further find that excision proceeds independently of target-site complementarity, relying strictly on donor-arm recognition; we term this "half-match" recombination, because a substrate matching only half of the bRNA is sufficient. We also find half-match activity during insertion, both *in vitro* and in a published genome-editing experiment, where it accounts for approximately half of non-target insertion reads. Half-match recombination provides both a mechanistic explanation for off-target insertion and a framework for the rational design of high-fidelity bridge recombinases.

**Highlights:**

- IS621 excision products support a copy-and-paste model
- Donor-only (DBL–DBL) recognition is sufficient to drive excision without a target
- Half-match explains 47% of non-target-site insertions by programmed bRNAs

## Introduction

IS110-family RNA-guided recombinases are a recently developed class of compact, programmable genome-editing tools.^1,2^ These systems use a bridge RNA (bRNA) coupled with the recombinase to programmably bring together a target and donor DNA and drive sequence-specific DNA insertion, excision, and inversion. Crucially, this occurs without generating double-strand breaks, in contrast to nuclease-based editors that depend on host double-strand-break repair.^3^ This capability provides a modular, compact approach to genome engineering, complementing CRISPR-Cas technologies and offering a solution to the persistent challenges of large-scale, scarless DNA integration.^1,4,5^

Current mechanistic understanding of these systems depends largely on a “full-match” model of insertion and excision. The two loops of the bRNA, target-binding loop (TBL) and donor-binding loop (DBL), bind fully complementary target and donor DNA to form a synaptic complex that drives double-stranded recombination during insertion.^2^ Similarly, excision is proposed to reverse this reaction through full TBL–DBL binding and double-stranded crossover, implying a cut-and-paste mechanism for IS621.^6^ Both reactions proceed through enzymatic cleavage and strand exchange at the central dinucleotide core.^2,6^ Notably, the related IS110-family member CazIS110 appears to use a distinct copy-and-paste mechanism for excision.^7^ These observations appear to conflict.^8^

Determining whether the IS110 circular intermediate is single- or double-stranded is critical to distinguishing between cut-and-paste and copy-and-paste mechanisms, yet this fundamental mechanism remains unresolved. Circularization represents a key intermediate stage in transposition, where recognition of the transposon’s flanking sequences (LT–RD and LD–RT) facilitates the formation of an LD–RD junction recognizable by the donor loop of the bRNA. If circularization proceeds via a double-stranded recombination mechanism, the excision reaction affects a direct transfer of the transposon without replication, consistent with a cut-and-paste pathway.^9^ Alternatively, single-stranded circularization followed by insertion leaves a complete ssDNA template for the transposon at the donor site, thereby allowing a copy-and-paste cycle.^9^ Although circularization has been documented for several IS110 elements,^10–15^ whether the excised intermediate is single- or double-stranded has not been directly established.^7^ In the IS621 TBL–DBL excision complex, top-strand cleavage and exchange are observed, whereas cleavage and exchange of the bottom strand have not been captured.^6^ Neither do structural insights preclude the possibility of single-stranded excision. Hiraizumi et al. detected rejoined LT–RT arms as evidence for double-stranded excision,^6^ but without a comparison to LD–RD formation efficiency it remains unclear whether LT–RT is the dominant product or a minor one.^6^ Therefore, whether IS621 excision proceeds through a single-stranded copy-and-paste route is still an open question in the field.

A second question is whether full bRNA recognition is required to initiate the reaction at all. The existing models for excision and insertion predict that both donor and target bRNA loops are fully bound to DNA within a single synaptic complex, so productive events should be confined to loci matching the complete bRNA sequence.^2,6^ Yet in published *Escherichia coli* genome-editing data,^1^ a fraction of off-target insertion events cannot be accounted for by the known full-length guide, and the enzymatic activity responsible for them remains unknown. Further, in an *in vitro* assay performed with IS621, recombinant product was generated when one donor substrate contained only the LD sequence rather than the full LD–RD match.^2^ A separate study from our lab identified large IS110 family transposons expanding their size by insertions at sites that are only complementary to half the bRNA guide.^16^ Whether partial bRNA recognition is sufficient to trigger recombination, and if so, whether the same pathway is a major driver of excision of the native transposon, where full complementarity is absent, has not been tested.

Thirdly, current understanding of the excision complex depends on a TBL–DBL architecture of synaptic complex; however, other results suggest a DBL- DBL-mediated excision process is possible. In a DBL–DBL reaction, two bRNA molecules would each contribute one DBL, rather than one bRNA contributing a TBL and another a DBL. For IS621, a DBL–DBL configuration has been modeled by superimposing the experimentally determined IS621–DBL–dDNA half-complex, an arrangement expected to recombine LD–RD-containing sequences without engaging LT–RT.^2^ IS621 supplied with a split bRNA that provides only the DBL can also form a DBL–DBL complex which, *in vitro*, catalyzes excision approximately 4.4-fold more efficiently than the wild-type TBL–DBL configuration (0.66% vs. 0.15%), suggesting that it could drive efficient excision in a native context.^6^ Current understanding of DBL–DBL recombination mostly depends on split bRNAs,^6,17^ but no study has directly compared the excision efficiencies of the TBL–DBL and DBL–DBL reactions using wild-type bRNA, nor is it clear how DBL–DBL recombination contributes to insertion and to off-target frequency. Resolving these questions is important for the rational use of these genome-editing tools.

In this study, we find that recognition of a partially complementary sequence by the bRNA, or “half-matches”, drives efficient excision and many of the off-target recombinations during insertion. Using *in vitro*-synthesized IS621 recombinase, we reconstituted *in vitro* recombination and found asymmetric rejoining of the excision products, with the circular single-stranded excision product forming ∼34-fold more efficiently than resealing of the transposon-flanking site, suggesting a copy-and-paste mechanism. Furthermore, we found that the excision reaction is not inhibited by mismatches or complete absence of target sites on the flanking region, showing target sequence independence. Re-analysis of genome-editing datasets showed that half-match events are present among non-target-site insertions, accounting for 47% of non-target-site insertion reads in that dataset. Our work provides insight into how excision occurs naturally and how to prevent off-target editing events.

## Results

### *In vitro* transcription and translation (IVTT) system produces active IS621 recombinase and bRNA

To investigate the mechanism of IS621 recombination (Fig. 1A), we used a reconstituted *in vitro* transcription–translation (IVTT) system to produce the active IS621 recombinase^18^ (via coupled transcription and translation) and its associated bRNA (via transcription alone)(Fig. 1B). This chemically defined, cell-free approach bypasses the challenges of protein purification and provides a controlled setting in which to probe the excision–insertion life cycle of IS621.

**Fig. 1.**
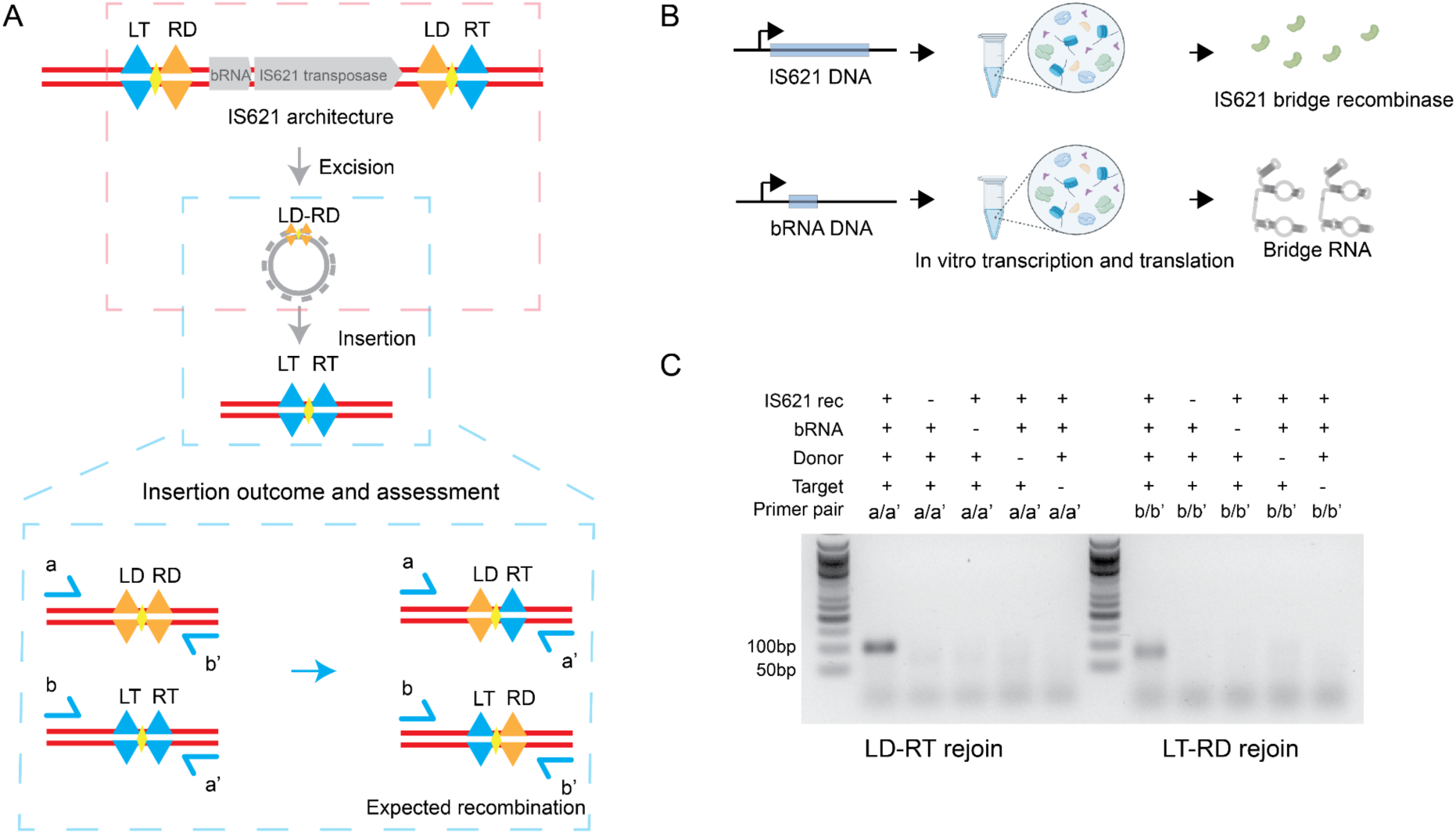
Mechanistic investigation of IS621 recombination using an *in vitro* transcription-translation (IVTT) system. **(A)** Schematic of the IS621 transposition process illustrating the two-step recombination process: excision (red) and insertion (blue). The excision process circularizes the donor sequence (LD–RD), which is subsequently recognized and inserted into the target site (LT–RT). Insertion recombination assay was used to validate the activity of IS621 recombinase and bRNA after *in vitro* synthesis. The canonical model depicts dual-strand crossover of two substrates, resulting in two recombinant products. **(B)** Overview of the cell-free IVTT setup used for the expression of the wild-type IS621 recombinase and its associated bridge RNA (bRNA). **(C)** Validation of recombination activity. Junction PCR was performed using specific primer sets (a/a’ and b/b’) to detect the recombinant products LD–RT and LT–RD, respectively. The gel analysis demonstrates efficient recombination, indicated by the presence of a recombinant band only in the presence of all required components. A representative gel from 3 independent experiments is shown.

We validated the activity of IS621 produced via the IVTT system using an existing PCR-based assay^2^. This assay uses the IS621 wild-type donor (LD–RD) and target (LT–RT) junction sequences as substrates to generate the predicted recombinant products, LD–RT and LT–RD. We adopted the LT–RT and LD–RD sequences and primer sets previously described with some modifications (Supplementary Table 1).^1,2^ Following IVTT, IS621, bRNA, donor DNA, and target DNA were combined (Fig. 1C) and recombination events were validated via junction PCR. Gel analysis confirmed that the recombinant band was generated only in the group containing active recombinase, bRNA, donor, and target sequences. In addition, one-pot transcription, translation, and recombination were also achieved (Fig. S1). Finally, Sanger sequencing verified the successful formation of the LD–RT and LT–RD sequences. Cell-free expression therefore yields active, specific IS621 and bRNA, giving us a defined system in which to dissect excision.

### IS621 excision products support a copy-and-paste model

An IS621 insertion leaves the element flanked by hybrid junctions: LT–RD on the left and LD– RT on the right (Fig. 2A). We used these two ends as the substrate for *in vitro* excision, which should regenerate LD–RD and LT–RT upon recombination (Fig. 2B). We detected the expected LD–RD junction but qualitatively observed far less LT–RT product (Fig. 2C). No recombinant band was detected in the negative controls (catalytically inactive IS621 or reactions lacking bRNA or donor). This result indicates asymmetric rejoining efficiency between LD–RD and LT– RT products.

**Fig. 2.**
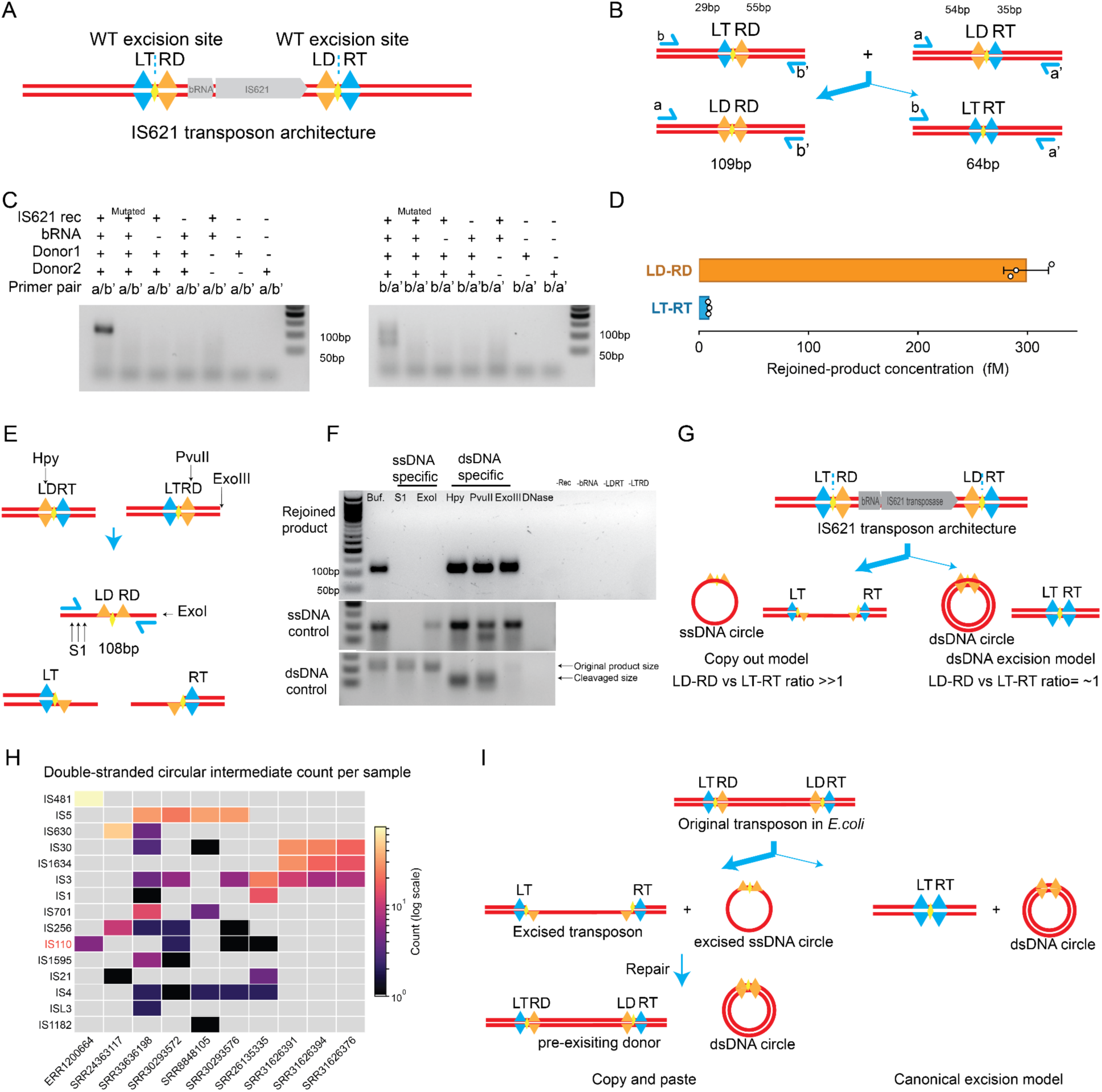
IS621 catalyzes asymmetric donor rejoining and generates single-stranded LD–RD junction products. **(A)** IS621 transposon architecture. The IS621 recombinase and its bridge RNA (bRNA) are flanked by two wild-type excision sites, LT–RD and LD–RT (dashed line, CT core; blue and orange symbols, target- and donor-side half-sites). **(B)** *In vitro* recombination assay design. Synthetic substrates spanning the LT–RD and LD–RT junctions were combined; recombination yields the LD–RD (109 bp) and LT–RT (64 bp) products, detected by junction PCR with primer pairs a/a′ and b/b′ (substrate arm lengths indicated). **(C)** Asymmetric substrate recognition. Recombination assays with the natural IS621 flanking sequences show efficient rejoining of the LD–RD junction (primers a/b′, left) but little LT–RT product (primers b/a′, right). Products require active IS621, bRNA, and both donors, and are lost with catalytically inactive (mutated) IS621 or upon component dropout. A representative gel from 3 independent experiments is shown. **(D)** Quantification of the rejoined products in (C). The LD–RD product is ∼34-fold more abundant than LT–RT, indicating asymmetric strand rejoining. Quantitative PCR was performed in triplicate; bars, mean ± s.e.m. **(E)** Schematic of the differential nuclease-sensitivity assay, showing the dsDNA-specific restriction sites (HpyCH4III, PvuII) and the S1/ExoI/ExoIII nucleases used to probe the LD–RD product. **(F)** Differential nuclease digestion of the rejoined product, an ssDNA control, and a dsDNA control with ssDNA-specific (S1, ExoI) and dsDNA-specific (HpyCH4III, PvuII, ExoIII) nucleases; the rejoined product is abolished by ssDNA-specific nucleases and resistant to dsDNA-specific nucleases, indicating a single-stranded product. A representative gel from 3 independent experiments is shown. **(G)** Two models for IS621 excision and their predicted junction stoichiometry. Left: single-strand cleavage and rejoining yields a single-stranded (ssDNA) circle (copy-out model), leaving the LT/RT sites largely intact, so LD–RD products exceed LT–RT products. Right: double-strand cleavage and recombination yield a double-stranded (dsDNA) circle (cut-out/excision model), producing LD–RD and LT–RT junctions at ∼1:1. The asymmetric rejoining in (D) is consistent with the copy-out model. **(H)** Detection of dsDNA circular intermediates *in vivo*. Publicly available nanopore datasets were mined for transposon tail–head junction reads (a pattern of circularization); the heatmap shows per-sample counts across IS families (IS110 highlighted), consistent with dsDNA circular intermediates for multiple IS elements. **(I)** Proposed model of IS621-mediated transposition. From the original transposon in *E. coli,* the copy- and-paste route (left) proceeds through single-strand excision to an ssDNA circle that host repair converts to a dsDNA circle, leaving the donor locus intact; a cut-and-paste route (right) excises a dsDNA circle and leaves an LT–RT scar. Our data, asymmetric rejoining and an ssDNA intermediate, support the copy-and-paste route.

To quantify LD–RD and LT–RT production in an *in vitro* excision reaction, we used quantitative PCR (qPCR) to detect recombinant products. Results indicate LD–RD production is ∼34-fold higher than LT–RT production (Fig. 2D), in accordance with the gel result (Fig. 2C). This indicates that LD–RD and LT–RT are not formed in the 1:1 stoichiometry expected of a cut-and-paste mechanism, but rather are consistent with a dominant copy-and-paste mechanism in which one strand is excised to form the circular transfer intermediate while the other maintains the intact donor sequence and a template for repair (Fig. 2G).

To further probe the single-stranded nature of excision, we performed nuclease-sensitivity assays that would differentiate ssDNA and dsDNA(Fig. 2E, F). We engineered dsDNA-specific restriction sites into the LD–RT (HpyCH4III) and LT–RD (PvuII) sequences as double-stranded probes and treated products with ssDNA-specific nucleases (S1, ExoI) and dsDNA-specific nucleases (HpyCH4III, PvuII, ExoIII). The *in vitro* product was digested by ssDNA-specific nucleases but resistant to dsDNA-specific nucleases (Fig. 2F). Matching the ssDNA controls, these results support that excision yields a single-stranded product and leaves the donor with single stranded gap.

We next confirmed that the excised molecule forms a covalently closed circle *in vivo*. Previous work has reported circular intermediates for IS110-family elements by junction PCR.^10–15^ Here we used a cell-based excision assay, where the IS621 and bRNA cassette was placed between the natural LT–RD and LD–RT flanking sites, so that the predicted excision product is a 2,579- bp circle (Fig. S2). Random-primed phi29 amplification, which generates tandem concatemers only from covalently continuous circular templates, yielded LD–RD junction concatemers with active IS621 in the presence of the donor-binding sites, but not with catalytically inactive IS621 or when it was absent. The monomer length (median 2,579 bp) matched the expected excised sequence and was distinct from the full-length plasmid (4,638 bp), ruling out whole-plasmid or linear background (Fig. S2C). The excised species is therefore a covalently closed circle carrying the LD–RD junction, rather than a nicked molecule or a protein-held synaptic complex. Searching publicly available nanopore whole-genome sequencing datasets, we recovered tail– head junction reads for IS110 and other IS elements, including IS481, IS5, IS630, IS1634, IS701, IS1595, IS4, and IS1182 (Fig. 2H), indicating that these junctions also form at native loci.

Based on our *in vitro* and *in vivo* findings, we propose a unified model for the IS621 transposition process (Fig. 2I). Guided by the bRNA, IS621 recognizes the left and right flanking sequences and performs single-strand cleavage, generating a circular ssDNA intermediate that carries the LD–RD junction and leaving an ssDNA donor locus. Host repair machinery then likely converts a fraction of the ssDNA circles to dsDNA circles, while the donor locus is repaired; the dsDNA circles are subsequently inserted at new target sites. This model classifies IS621 as a copy-and-paste system as strand cleavage and repair effectively duplicate the transposon, while the subsequent insertion follows a cut-and-paste mechanism,^2^ consistent with the copy-and-paste model reported for CazIS110.^7,8^

### Donor half-site recognition is sufficient for efficient excision

Since our data show strong bias towards LD-RD rejoining over that of LT-RT, we hypothesized that the LD and RD sites, rather than a full match (LT–RD and LD–RT), are sufficient to drive excision. To test whether the target halves of the flanking sequences are required for excision, we repeated our *in vitro* recombination assay using the substrate design of Fig. 2B but with the target sites mutated: the LT and RT sequences were altered to abolish bRNA complementarity while the donor arms were left intact. Rejoining of the LD–RD junction remained efficient and was confirmed by Sanger sequencing (Fig. 3A). Excision therefore does not require an intact target site, indicating that donor-side recognition alone is sufficient to assemble a productive complex.

**Fig. 3.**
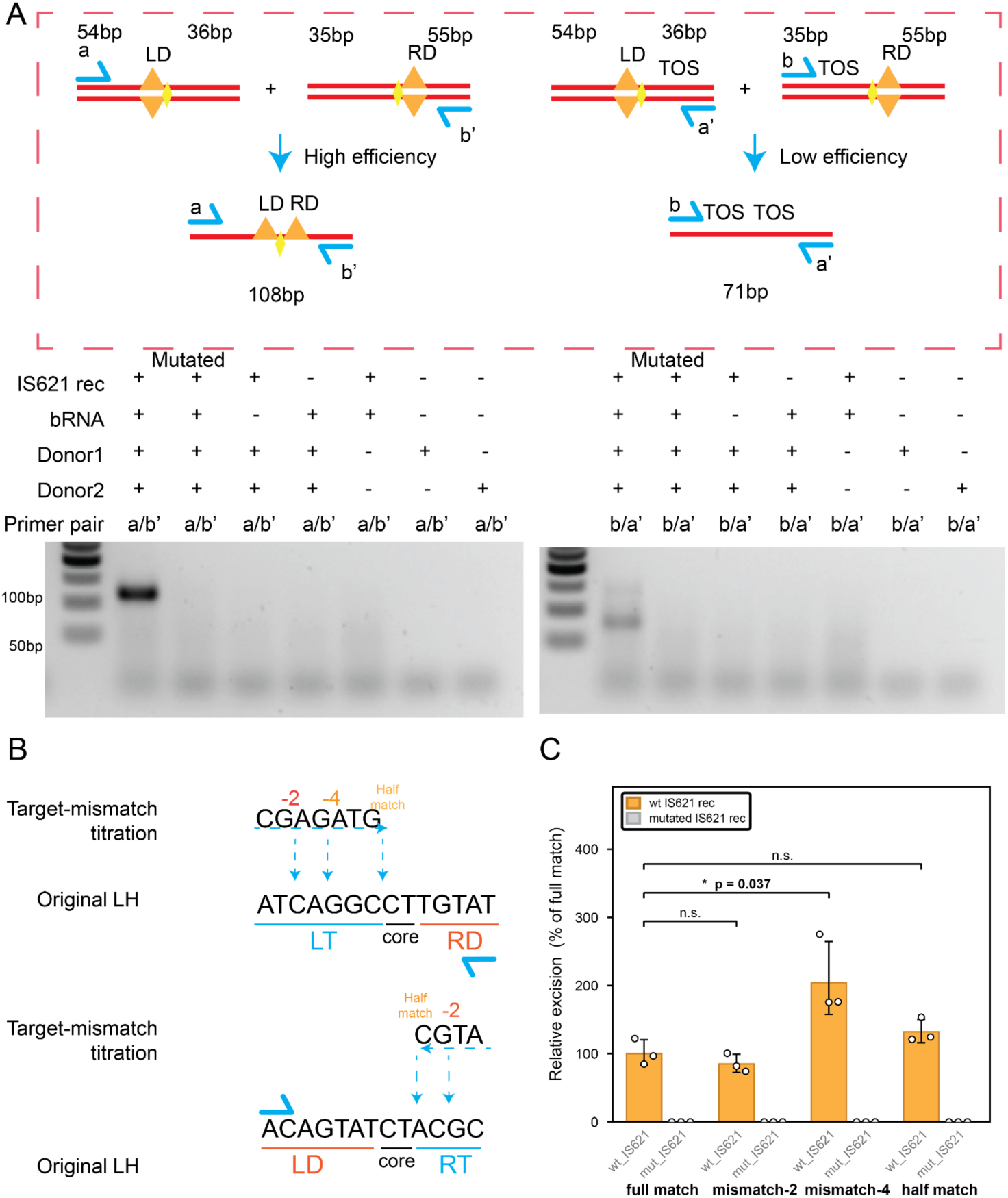
Half-match sufficiency of driving excision. **(A)** Donor-site sufficiency for recombination. Substrates of the same design as in Fig. 2B, but carrying mutated target (LT and RT) sequences that abolish bRNA complementarity, still support efficient LD–RD rejoining. Products were validated via junction PCR and confirmed by Sanger sequencing; no recombinant band was detected with catalytically inactive IS621 or on omission of the recombinase, bRNA, or donor. Formation of the other-strand (TOS) rejoining product is shown on the right. A representative gel from 3 independent experiments is shown. **(B)** Target-site mismatch titration reveals that LD–RD rejoining does not require target-site recognition. Mismatches were introduced into the LT and RT sequences pairwise from the outer edge toward the CT core, leaving the donor arms (LD and RD) intact; −2 and −4 indicate the number of mutated nucleotides per arm, and “half match” denotes a target arm with no remaining complementarity. **(C)** Quantification of LD–RD excision product (LD–RD qPCR). Bars show the geometric mean with individual replicates overlaid (n = 3 biological replicates, each normalized to full match); error bars, geometric SD. Comparisons to full match were made by unpaired two-tailed Student’s t-test on the calculated concentrations; *p < 0.05; n.s., not significant.

To quantify how much target complementarity contributes to excision efficiency, we titrated target-site mismatches and measured excision by qPCR (Fig. 3B, C). Starting from the wild-type target sequence, we introduced mismatches pairwise to progressively abolish bRNA base-pairing from the edge toward the core, and used a junction primer amplifying across the LD–RD junction to measure product abundance after *in vitro* recombination. This allowed a direct comparison of excision efficiency across the mismatch series. Introducing target-site mismatches did not measurably reduce excision efficiency. Interestingly in the Mismatch-4 group there was a significant increase of the rejoined product, compared with full match substrates. Within the current full-match excision model, this is unexpected, and it suggests that IS621 possesses an alternative excision pathway that efficiently rejoins the donor without target recognition, potentially without TBL engagement. Given that a DBL–DBL complex has been proposed structurally and appears as a prominent off-target configuration in human genome-editing data, and that target mismatching did not reduce the LD–RD product, we hypothesize that this activity reflects DBL–DBL recombination.

### Non-specific IS621 insertions result from half-match recombination

Because excision proceeded with only two half-site matches to donor sequences, we asked whether incomplete matching also permits insertion, thereby allowing the recombinase to act on partially matched loci across the genome. We synthesized a DNA donor sequence containing the normal LD-RD bRNA recognition site and a range of target sequences with various bRNA binding motifs. Next, we assessed their activity using IVTT-based recombination assays. Surprisingly, we observed that when one substrate harbored the native LD–RD junction, the presence of a single complementary “half-site” (new LD, nRD, nLT, or nRT) on the partner DNA was sufficient to trigger efficient recombination, yielding detectable recombinant products. These products required catalytically active IS621, because no amplicon was detected with mutated (inactive) IS621, or when the recombinase, bRNA, or partner substrate was omitted (Fig. 4A), excluding PCR template switching between the co-present substrates. These results broadened our definition of a half-match to any bRNA loop and substrate pair that are complimentary across the CT core and one flanking arm, while the arm on the opposite side of the core is not complementary. This one-sided match is in contrast to the canonical reaction, which requires both arms of both loops to pair before catalysis.

**Fig. 4.**
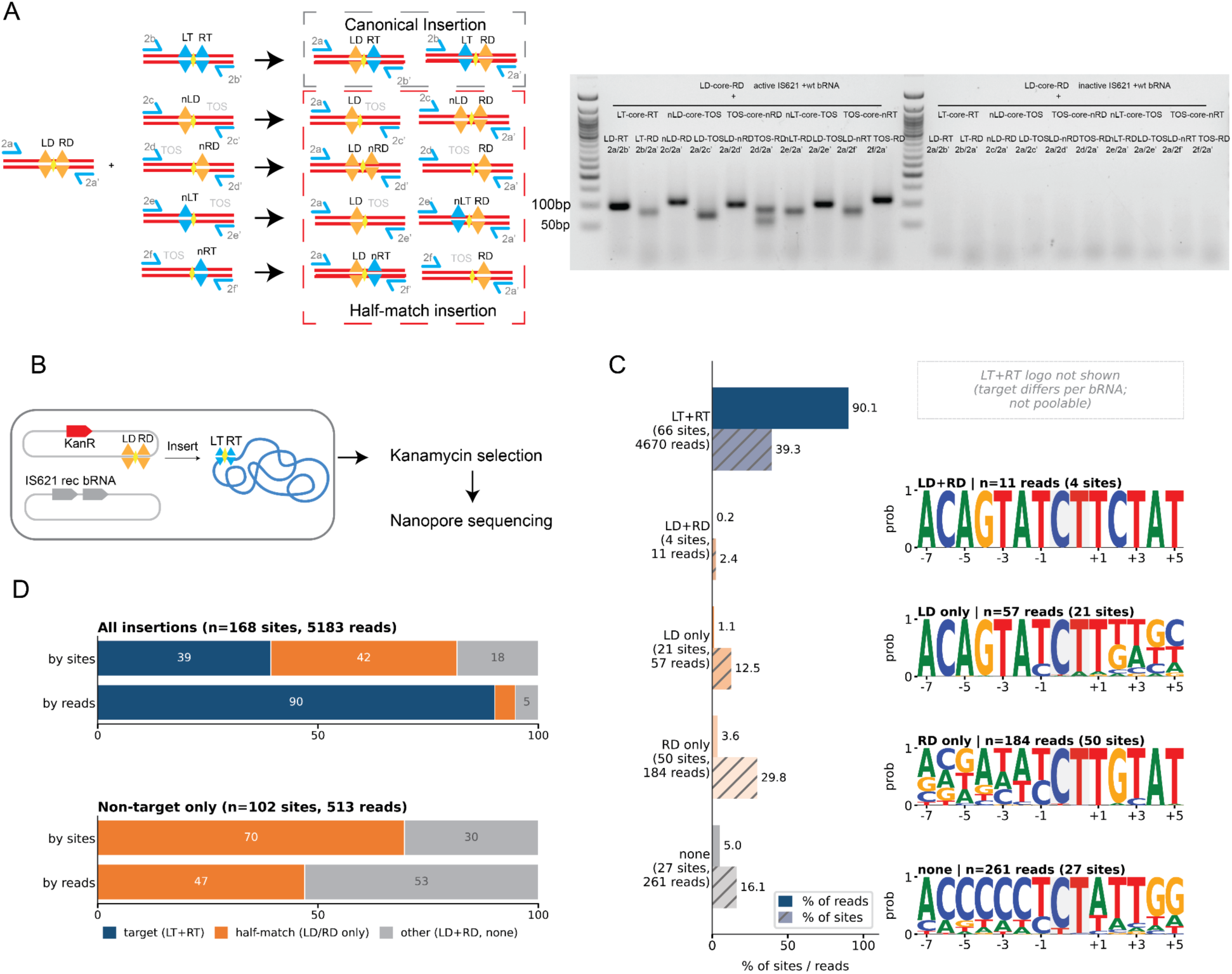
Half-match recombination bypasses the full-match requirement and explains off-target insertion events. **(A)** Half-match sufficiency for insertion recombination. Recombination assays using synthetic substrates with single complementary “half-site” motifs (nLD, nRD, nLT, or nRT) and the other strand (TOS) with no complementarity to the guide. Recombination products are detected across all tested combinations, indicating that partial guide engagement within the reaction core is sufficient to trigger productive recombination. No products were detected with catalytically inactive IS621 or when the recombinase, bRNA, or partner substrate was omitted. **(B)** Schematic of the *in vivo* insertion assay. Overview of the whole-genome sequencing (WGS) approach used to profile and map insertion sites for both programmed and wild-type (WT) bRNAs. The dataset was previously described.^1^ **(C)** Recognition categories of recovered insertion sites. Left: percentage of insertion reads (solid) and of insertion sites (hatched) assigned to each category in the programmed dataset. LT+RT (66 sites, 4,670 reads), LD+RD (4 sites, 11 reads), LD-only (21 sites, 57 reads), RD-only (50 sites, 184 reads) and none (27 sites, 261 reads). Right: sequence logos of the 14-bp recombination windows recovered in each category; the LT+RT logo is not shown because the programmed target sequence differs between bRNAs and is therefore not poolable. **(D)** Composition of insertion events. Top: all insertions in the programmed dataset (n = 168 sites, 5,183 reads). Bottom: non-target-site insertions only, within the programmed dataset (n = 102 sites, 513 reads), of which half-match events (LD-only + RD-only) account for 47% of reads and 70% of sites. Half-match events correspond to 4.7% of all reads in the programmed dataset and ∼1.9% across the full dataset (n = 341 sites, 13,169 reads); programmed bRNAs show a higher frequency of half-match insertion than WT bRNAs (Fig. S3, S7).

To investigate the relevance of this half-match activity to IS110’s use as an editor, we re- analyzed previously published IS621 editing outcomes. These data, from Durrant et al., profile genome-wide insertion sites for the native bridge RNA alongside four reprogrammed bridge RNAs, giving a direct comparison of insertion specificity between natural and engineered targeting.^1^ We classified insertion sites into discrete categories: LT+RT (canonical target-like), LD+RD, LD-only, RD-only, and none (Fig. 4B, C, Supplementary Table 2). Because the native right target guide (RTG) is only 4 bp long, separating LT- and RT-only from LT+RT can not be done confidently, so LT-only and LT+RT containing sequences were grouped into the LT+RT bin, and RT-only is excluded from categorization. Our analysis revealed that while 90.1% of programmed insertions targeted the canonical LT–RT junction, there was a frequent occurrence of “half-match” insertion events, where the insertion site matched only one half of the full-guide sequence (LD-only or RD-only). Excluding LT–RT insertion events, donor half-match sites, matching one arm of the donor recognition sequence, captured 47% of non-target-site insertion reads across the programmed bridge RNAs (Fig. 4D; equivalent to 4.7% of all insertion reads in this dataset; n = 168 sites, 5,183 reads), a 48-fold enrichment over the expectation for random insertion at genomic CT core sites (permutation test relocating each insertion event to a random CT site; p = 1.8 × 10⁻⁸). Specifically, LD-only and RD-only events accounted for 1.1% and 3.6% of total insertion reads, respectively (corresponding to 12.5% and 29.8% per site). Interestingly, RTG-extended bRNAs showed fewer half-match insertions (Fig. S6) and programmed bRNAs exhibited a higher propensity for half-match insertion compared to wild-type bRNAs (Fig. S7). Quantitatively, half-match-mediated recombination accounted for approximately 1.9% of all insertion reads across the full dataset (n = 341 sites, 13,169 reads; 23% of insertion sites), and explained 67.1% of non-target-site insertion reads for the native RTG bRNA2 (116/173 reads; Fig. S4, S5). These data suggest that the “off-target” propensity of bridge recombinases is an intrinsic feature of their ability to recombine at half-matches.

## Discussion

Our work provides evidence that IS621 follows a copy-and-paste transposition process initiated by half-matches between the bRNA and the LD and RD sequences flanking the insertion sequence. We find that excision’s dominant mechanism peels off a single-stranded intermediate, leaving the other strand of the donor locus intact, a signature of copying rather than cutting (Fig. 2). This updates from previous description of IS621 as a cut-and-paste element,^2,6^ instead placing it alongside the copy-and-paste mechanism of CazIS110.^7,8^ Furthermore, excision proceeds efficiently with matches to LD and RD sequences even when the LT and RT are absent altogether (Fig. 3). The same half-match mechanism that drives native excision also explains many of the off-target insertion products of programmed editing (Fig. 4D). Together these findings reframe the mechanism behind a promising new class of programmable editors, relevant both to their natural lifecycle and use in synthetic biology.

The strandedness of the circular intermediate is central to understanding the mechanism, and to predicting the outcomes of genetic engineering. Although IS110 circularization has been examined before,^10–15^ direct evidence for whether the post-cleavage intermediate is single- or double-stranded has been limited. Notably, the excision structure reported by Hiraizumi et al. focuses on top-strand cleavage and top-strand exchange.^6^ That study also reported repair of the LT–RT junction after excision, indicating dsDNA cleavage consistent with a cut-and-paste mechanism. However, our results show that LD–RD rejoining is highly favorable compared to LT–RT rejoining *in vitro* (Fig. 2C, D). Consistent with only a single strand being removed during excision, our nuclease treatment assay supports the circular excision intermediate consisting of ssDNA (Fig. 2E, F). Together, these data support a model in which IS621 preferentially cleaves and rejoins a single strand to generate a ssDNA LD–RD excision product that is subsequently resolved into a circular intermediate during copy-out transposition.

This ssDNA excision has unexpected independence from the LT-RT target sequences. Titrating mismatches into the target site did not reduce excision efficiency (Fig. 3C), so the reaction is not aided by the target sequence, implying a synaptic complex in which two bRNAs each contribute a DBL. Donor-only activity has been noted before, but as an anomaly rather than a mechanism.^2,2,6^ In addition, DBL–DBL products were identified as off-target events in IScro4 editing of human cells.^17^ Our data indicate that this configuration operates on the native element with wild-type bRNA and is an efficient route for excision. Why donor-only complexes should in some cases excise more efficiently remains open. One candidate is the handshake guide, a short pairing element that biases the wild-type complex toward insertion and therefore resists the reverse reaction, excision. Because a DBL–DBL complex carries the same donor-side element on both loops, this built-in directional preference is absent, which could let excision proceed more freely.^6^

Donor-only excisions appears to be one instance of a more general property in which IS621 recombination can be initiated by a half-match mechanism. The Donor-side half-match has similarities to the direct strand-transfer activity reported for CazIS110,^7^ which does not require full complementarity; but our observation expands this activity beyond TBL–DBL formation and CazIS110. Half-match-mediated insertions have also been observed in a bioinformatic survey of how IS110 family transposons expand.^16^ Our observation, in the context of literature observations, suggests that half-match recombination occurs throughout the IS110 family.

This half-match promiscuity combined with the copy-and-paste mechanism may confer an evolutionary advantage on IS110-family transposons. Unlike the canonical reaction, which relies on 11 bp (native) or 14 bp (with extended RTG) of recognition, half-match recombination can proceed from a single site as short as the 5-bp RD motif (Fig. 4A). This lowered threshold along with the copy-and-paste mechanism could facilitate propagation and may help explain the high copy numbers of IS110 elements, reaching 106 copies per genome in *Coxiella burnetii* strain NL3262.^19^

The half-match pathway also provides a mechanistic explanation for a substantial fraction of off-target events in bridge-recombinase editing, arguing for a shift toward rational bRNA design that disfavors half-match recombination while preserving canonical insertion. Consistent with this, extending the right target guide (RTG) improved specificity by reducing half-match events (Fig. S6). Interestingly, wild-type bRNAs showed a lower half-match propensity than reprogrammed bRNAs. Whether this reflects sequence features that suppress off-target excision or an incidental property of the native guides cannot be resolved here and is an important next step toward high-fidelity editors.

In summary, we identify half-match recombination as an alternative IS621 pathway that can proceed without canonical target complementarity, is associated with a copy-and-paste mechanism, and offers a mechanistic explanation for a substantial class of off-target insertion events. This framework clarifies the IS110 transposition process and offers a rational route toward high-fidelity genome editors that decouple therapeutic insertion from the ancestral excision pathway.

## Methods

### *In vitro* transcription and translation (IVTT) of IS621 recombinase and bridge RNA

The IS621 recombinase and bridge RNA (bRNA) were synthesized utilizing the PURExpress *In Vitro* Protein Synthesis Kit (New England Biolabs, E6800S). For each expression reaction, 200 ng of template DNA was added to the IVTT mixture, which was supplemented with RNase inhibitor (New England Biolabs, M0314S) according to the manufacturer’s recommendations. The DNA expression templates were engineered to comprise a T7 promoter, the respective target coding sequence (IS621 or bRNA), and a T7 terminator. The assembled reactions were incubated at 30 °C for 2.5 hours. To serve as a negative control, a catalytically inactive IS621 mutant was generated and expressed, in which all putative active site residues (the DEDD motif and the catalytic serine) were substituted with alanine. Following the incubation, the reaction mixtures were immediately transferred to -80 °C for storage until downstream applications.

### *In vitro* Recombination (IVR) assay

The *in vitro* recombination reactions were performed in a 20 μL total reaction volume. The reaction mixture consisted of 1X T4 DNA Ligase buffer (New England Biolabs, B0202S), 200 mM NaCl, double-stranded DNA donor substrates (e.g., LD–RD and LT–RT), and 0.8 μL of RNase inhibitor. To initiate the reaction, 1 μL of unpurified guide RNA (bRNA) and 1 μL of the IS621 protein (wild type or mutant), both previously generated via cell-free transcription-translation systems (incubated at 30 °C for 2.5 hours), were added to the mixture.

The assembled reactions were incubated at 30 °C for 1.5 hours. Following the incubation, the reactions were temporarily held at 4 °C for immediate downstream analysis or stored at -80 °C. The rejoining products and specific junction events were subsequently detected and amplified utilizing junction PCR.

### Nuclease digestion assay

To characterize the strand structure and integrity of the DNA intermediates or rejoining products, nuclease digestion assays were performed using a panel of structure-specific endonucleases and exonucleases, including S1 Nuclease (Thermo Scientific, EN0321), Exonuclease I (New England Biolabs, M0293S), Exonuclease III (New England Biolabs, M0206S), HpyCH4III (New England Biolabs, R0618S), PvuII (New England Biolabs, R0151S) and DNase I (New England Biolabs, M0303S). Digestion reactions were assembled in a 10 μL total volume, comprising the corresponding 1X reaction buffer, 5 U of the designated nuclease, and the target DNA substrate (e.g., *in vitro* rejoining products, single-stranded DNA, or double-stranded DNA controls). For nucleases requiring lower working concentrations (e.g., S1 Nuclease and Exonuclease III), enzymes were serially diluted in their respective 1X buffers prior to addition.

The assembled reaction mixtures were incubated at 30 °C for 1 hour, followed by immediate heat inactivation at 80 °C for 20 minutes (specific incubation temperatures, such as 37 °C for Exonuclease I, were adjusted according to the manufacturer’s guidelines). The resulting digested products were subsequently analyzed via direct 3% agarose gel electrophoresis or utilized as templates for downstream junction PCR detection.

Synthetic single-stranded DNA (ssDNA) and double-stranded DNA (dsDNA) controls mimicking the post-rejoining products were prepared and subjected to digestion. For the dsDNA control, the complementary top and bottom oligonucleotides were mixed at equimolar concentrations in 1X T4 DNA Ligase buffer. The mixture was heated to 95 °C for 5 minutes and allowed to gradually cool to room temperature over 50 minutes to facilitate proper annealing.

Prior to digestion, the ssDNA control substrates underwent a thermal pre-treatment to minimize the formation of secondary structures (e.g., 3′ hairpins or partial duplexes). Briefly, the ssDNA was incubated at 95 °C for 5 minutes and immediately snap-cooled on ice for 2 minutes.

The subsequent nuclease digestion reactions were performed in a 10 μL total volume containing the corresponding 1X reaction buffer, 5 U of the specific nuclease, and the pre-treated control substrates (2400 ng for ssDNA and 240 ng for dsDNA to ensure visualization). The reactions were incubated at 30 °C for 1 hour (37 °C for Exonuclease I), followed by immediate heat inactivation at 80 °C for 20 minutes. Digestion profiles were then directly analyzed utilizing 3% agarose gel electrophoresis.

### Target Mismatch Titration assay

To characterize if target sequences have impact on excision efficiency, mutations on the target sequence were introduced and served as donor substrates for in vitro recombination, followed by quantitative PCR to detect LD–RD rejoin production. In mismatched groups, mutations were introduced starting from the other edge towards the core sequence (CT). 2 mismatches, 4 mismatches and all target sequence mismatch (Half-match) groups were prepared according to sequence presentation in Fig. 2B. After preparation of cell free synthesis of wild-type IS621, mutated recombinases and wild-type bRNA as previously described, In vitro recombination reactions were carried out in a total volume of 10 µL. Each reaction mixture contained 1X T4 DNA ligase buffer (NEB), 200 mM NaCl, 0.4 µL RNase inhibitor, 0.5 µL of the TXTL-expressed bRNA, and 0.5 µL of the TXTL-expressed IS621 protein. Subsequently, 7.5 nM of each diluted DNA donor substrate (Donor 1 and Donor 2) was added to the mixture. The recombination reactions were incubated at 30°C for 1.5 hours to allow for junction formation, followed by a rapid cool-down and hold at 4°C. Reactions were either immediately quantified or stored at - 80°C.

The efficiency of donor rejoining and junction formation was evaluated via real-time quantitative PCR (qPCR) utilizing the SsoAdvanced Universal SYBR Green Supermix (Bio-Rad, 1725270). The reactions were prepared consisting of 1X SYBR Green Supermix, 1 µM of target-specific forward primer and reverse primer, and 0.5 µL of the unpurified IVR reaction serving as the template, and water was used to reach a 15 µL final volume. LD–RD (109 bp) amplicons were quantified against independent SYBR-Green standard curves prepared from a common 10-fold dilution series (5×10⁵ to 5 fM, 6 points, triplicate).

### Recombination-site framing and half-match classification

Genomic insertion sites were obtained from Supplementary Table 3 of Durrant et al. (2024), comprising insertion loci recovered for the wild-type IS621 bridge RNA (T-WT/D-WT) and for four reprogrammed bridge RNAs (Bridge RNA 1–4), each assayed with the native (4-bp) or extended (7-bp) right target guide (RTG).^1^ For each insertion locus, we used the 14-bp genomic window provided in the table, framed on the empirically defined recombination core such that positions 1–7 constitute the 7-bp left arm, positions 8–9 the central CT core, and positions 10– 14 the 5-bp right arm. The donor recognition sequence (DBL target) is 5′- ACAGTAT·CT·TGTAT-3′, defining a donor left arm (ACAGTAT) and a donor right arm (TGTAT); the target (TBL) recognition sequence is the bridge-RNA–specific programmed target given in the table.

Each locus was split at the core, and its left (7-bp) and right (5-bp) arms were independently compared, by Hamming distance in the fixed frame, to the corresponding arms of both the target (TBL) and donor (DBL) sequences. An arm was scored as “matched” when it differed from the reference arm by ≤1 mismatch. Loci were then assigned to one of five recognition categories: LT+RT (either target arm matched, i.e., target-recognized/on-target-like), LD+RD (both arms matched the donor), LD-only (left arm matched the donor only), RD-only (right arm matched the donor only), and none (no arm matched either reference). Because the native RTG is only 4 bp, a genuine LT+RT locus is frequently scored as LT-only when the short right arm falls below the match threshold; we therefore did not treat LT-only as a separate class and analysed LT-only together with LT+RT as a single target-recognized category. Sites with a single donor-arm match (LD-only and RD-only) were defined as donor half-match sites. We verified that the fixed-frame arm distances reconstitute the authors’ reported 11-bp Levenshtein specificity distances, and confirmed that match/no-match calls were unchanged when Hamming distance was replaced by Levenshtein distance (indel-tolerant alignment affected only loci already scored as non-matching).

### *In Vivo* Circle Enrichment and Rolling Circle Amplification (RCA)

To investigate the *in vivo* generation of circularized DNA intermediates by the IS621 system, an anhydrotetracycline (aTc)-inducible expression and targeted enrichment pipeline was established. *E. coli* BL21(DE3) strains harboring aTc-inducible expression cassettes for the IS621 recombinase and its cognate donor substrates were cultured to an optical density (OD) of 0.8. Protein expression and *in vivo* recombination were induced by the addition of aTc to a final concentration of 200 nM, followed by incubation for 15 hours at 30°C. Post-induction, cells were pelleted, and total plasmid DNA was extracted using standard column purification.

To selectively enrich newly formed circular episomal intermediates and eliminate background interference from parental plasmids and genomic DNA, the extracted nucleic acids were subjected to a two-step enzymatic depletion protocol. First, samples were digested with the I-SceI restriction endonuclease (New England Biolabs, R0694S) (37°C for 2 hours, followed by heat inactivation at 65°C for 20 min) to specifically linearize unreacted parental plasmids. The linearized products were subsequently treated with Plasmid-Safe™ ATP-Dependent DNase (LGC Biosearch Technologies, E3101K) (37°C for 2 hours, followed by 70°C for 30 min), a highly specific exonuclease that degrades linear double-stranded and single-stranded DNA while leaving closed-circular topologies intact. The enriched circular fraction was then purified utilizing a DNA clean-up kit.

The purified circular intermediates were subjected to Rolling Circle Amplification (RCA) to generate high-molecular-weight concatemers for sequencing. Reactions were assembled using Exo-Resistant Random Primer (Thermo Scientific, SO181) (final concentration 2.5 μM) and denatured at 95°C for 3 minutes to facilitate primer binding, followed by cooling to room temperature. Isothermal amplification was initiated by the addition of phi29 DNA Polymerase (10 U; New England Biolabs, M0269S) supplemented with recombinant albumin (0.1 mg/mL; New England Biolabs, B9200S) and 1 mM dNTPs (New England Biolabs, N0447S). The RCA reactions were incubated at 30°C for 12 hours to ensure robust signal amplification, followed by thermal inactivation of the polymerase at 65°C for 10 minutes. The resulting concatemeric products were directly subjected to long-read Nanopore sequencing to confirm the target sequence, junction architecture, and precise topological features of the circular intermediates.

### IS-family quantification and visualisation

Ten publicly available Oxford Nanopore long-read datasets were analysed as deposited, without re-basecalling; sequencer model and basecalling software are as reported in the original submissions.^20–23^, and in NCBI BioProject: PRJNA497605, PRJNA497605, PRJNA991976, PRJNA1264753. Insertion sequence (IS) elements were identified with ISFinder,^24^ which records for each element its sequence and length, assembly-graph coordinates and orientation, up to 80 nt of flanking sequence on either side, and junction-read counts; boundaries were resolved by local re-assembly of spanning reads or, where this failed, from the single longest supporting read, and downstream analyses did not condition on the class of boundary evidence. Because excision as a circular intermediate juxtaposes the 3′ ("tail") and 5′ ("head") ends of an element, reads spanning this junction in tail-to-head orientation were counted per element (n_TH) alongside the two insertion junctions (genome-to-head, tail-to-genome), and only elements with n_TH ≥ 1 were carried forward. To recognise the same element across samples despite long-read error, elements were clustered into IS systems with MMseqs2 at 95% nucleotide identity over 80% of the shorter sequence,^25^ in independent batches of ∼25 samples; clusters are therefore batch-scoped, the ten samples analysed here span batches 001–004, and all cross-sample comparisons are made at family rather than system level. Each cluster was annotated against ISfinder using blastn from BLAST+ against a database snapshot downloaded,^24,26^ retaining hits with E-value ≤ 1 × 10⁻¹⁰, ≥ 80% nucleotide identity and ≥ 80% query coverage, and was assigned its dominant family only if at least one member returned a hit; unmatched clusters were excluded and elements inherited their cluster’s family. Within each sample × family bin, sequences were collapsed by 15-mer Jaccard similarity from longest to shortest, retaining a sequence only if its similarity to every retained sequence was below 0.90 (≈99.6% nucleotide identity for sequences of comparable length,^27^ giving a 10 × 15 count matrix. The matrix was handled with NumPy,^28^ reordered on both axes by average-linkage hierarchical clustering of Euclidean distances ^29^ after adding a pseudocount of 0.1 for ordering purposes only, and rendered with Matplotlib ^30^ on a logarithmic colour scale (vmin = 1, reversed magma) with zero cells masked in grey and counts overlaid as text.

## Resource availability

### Lead contact

Further information and requests for resources and reagents should be directed to and will be fulfilled by the lead contact, Carlotta Ronda and Benjamin E. Rubin.

### Materials availability

All nucleic acid sequences used in this study can be found in Supplementary Table 1. All plasmids generated in this study are available from the lead contact on request.

### Data and code availability

Nanopore sequencing data generated in this study, from rolling-circle amplification of the in vivo excision products, have been deposited at the NCBI Sequence Read Archive and are publicly available as of the date of publication under BioProject accession PRJNA1522500. This study re-analyzed previously published data. Genomic insertion sites for the wild type and reprogrammed bridge RNAs were obtained from Supplementary Table 3 of Durrant et al. (2024). Codes related to the analysis were deposited at 10.6084/m9.figshare.33437989.The publicly available nanopore whole-genome sequencing datasets used for the tail–head junction survey are available under accessions ERR1200664, SRR8848105, SRR24363117, SRR26135335, SRR30293572, SRR30293576, SRR31626376, SRR31626391, SRR31626394 and SRR33636198. Any additional information required to reanalyze the data reported in this paper is available from the lead contact on request.

## Limitations of the study

While our biochemical and sequencing analyses provide evidence for half-match recombination, a structural basis for this mechanism remains to be established. Furthermore, our off-target analysis relies on the re-analysis of previously published datasets; as these experiments utilized antibiotic-based selection, they may not fully reflect the enzyme’s intrinsic recombination preference. Future investigations employing deep sequencing of non-selected edited cell populations will be necessary to decouple enzymatic specificity from potential selection bias.

## Author contributions

B.X., K.H., H.Y., J.R.P., B.E.R. and C.R. conceived and designed the experiments. B.X. and H.Y. performed the experiments. B.X., K.H., H.Y. and B.E.R. analyzed the data. B.X. wrote the paper with input from B.E.R., C.R., J.R.P., and K.H.

## Declaration of generative AI and AI-assisted technologies in the writing process

During the preparation of this work, the author(s) used Claude in order to write code for data analysis and writing improvement. After using this tool, the authors reviewed and edited the content as needed and take full responsibility for the content of the published article.

## Acknowledgements

We’d like to thank our funders. **JBEI:** This material is based upon work at the Joint BioEnergy Institute (JBEI) supported by the U.S. Department of Energy, Office of Science, Biological and Environmental Research Program under contract DE-AC02-05CH11231 with Lawrence Berkeley National Laboratory. **The Audacious Project:** This work was supported in part by Lyda Hill Philanthropies, Acton Family Giving, the Valhalla Foundation, Hastings/Quillin Fund— an advised fund of the Silicon Valley Community Foundation, the CH Foundation, Laura and Gary Lauder and Family, the Sea Grape Foundation, the Emerson Collective, Mike Schroepfer and Erin Hoffman Family Fund—an advised fund of Silicon Valley Community Foundation, and the Anne Wojcicki Foundation through The Audacious Project at the Innovative Genomics Institute. **Leona M. and Harry B. Helmsley Charitable Trust:** This work was funded in part by grant [G-2302-06692] from The Leona M. and Harry B. Helmsley Charitable Trust. **Shurl and Kay Curci Foundation:** This work was supported by a Research Award from the Shurl and Kay Curci Foundation (https://curcifoundation.org) to the Innovative Genomics Institute Genomic Tool Discovery Program at UC Berkeley, awarded to B.E.R. **Innovative Genomics Institute:** This work was supported in part by the Innovative Genomics Institute. **m-CAFEs:** This work was supported by m-CAFEs Microbial Community Analysis and Functional Evaluation in Soils, a Science Focus Area led by Lawrence Berkeley National Laboratory based upon work supported by the US Department of Energy, Office of Science, Office of Biological and Environmental Research under contract number DE-AC02-05CH11231. CR acknowledges funding support by Food Allergy Fund and SANDIA National Laboratory.

The authors thank Sophia Swartz, Leo Song, Jack Demaray, Zachary LaTurner, Siqi Yang, Brady Cress, Jonathan Martinson, and other members of the Rubin, Ronda, and Cress labs from the Innovative Genomics Institute for comments on improving the manuscript. The authors also thank Conner Langeberg and Kate Shulgina from the Innovative Genomics Institute for helpful discussions. The authors also thank the UC Berkeley DNA Sequencing Facility, especially Brian McCarthy, for help and discussions on DNA sequencing.

