## Supplementary Information for "Half-match recombination drives bridge RNA-guided excision and off-target insertion"

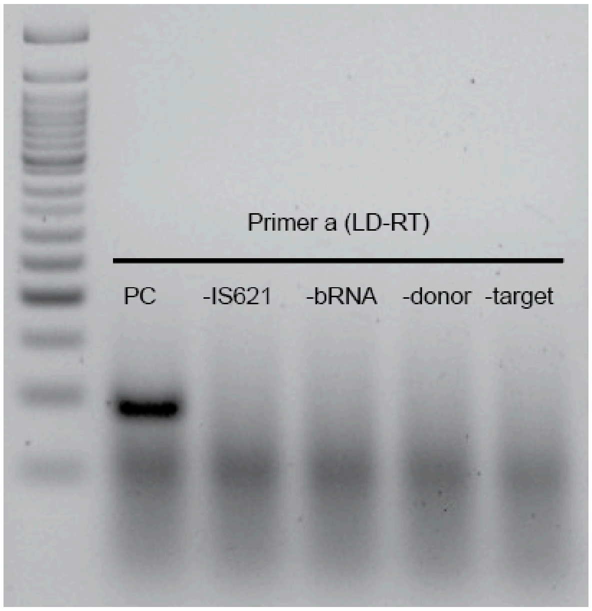


**Fig. S1. One-pot in vitro transcription, translation and recombination.**

The IS621 and bRNA expression templates, together with the donor and target sequences used in Fig. 1C, were combined in a single IVTT reaction to test whether expression and recombination can proceed in one pot. The recombinant LD–RT product was detected only when all required components were present.

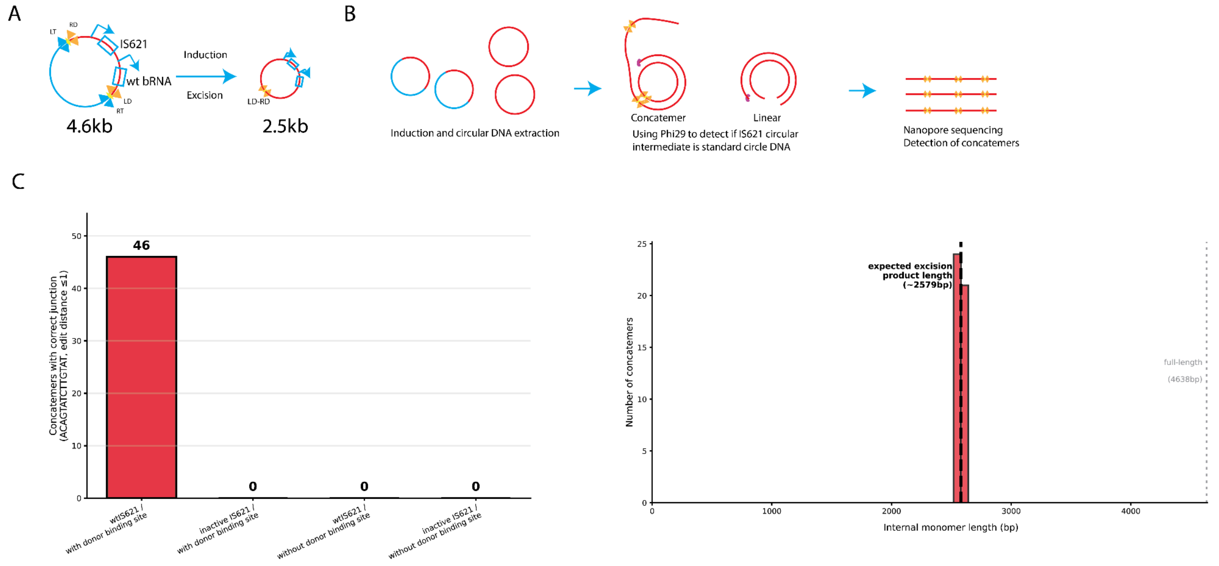


**Fig. S2. Rolling-circle amplification (RCA) confirms that the excised transposon intermediate is a covalently closed DNA circle.**

(A) Four plasmid constructs were built to assay circular products, combining the presence or absence of donor cleavage sites with catalytically active or inactive IS621. In the cleavage-site–containing constructs, the IS621 LT–RD and LD–RT sites flank the IS621 and bRNA expression cassette; upon excision, a 2.5-kb circular intermediate is released from the 4.6-kb parental plasmid.

(B) Phi29-based RCA followed by nanopore sequencing was used to test whether the excision product is covalently closed. Phi29 amplifies processively only on a covalently closed circular template, generating concatemers that are subsequently detected by nanopore sequencing.

(C) Concatemers were detected only for the construct carrying both the donor cleavage sites and active IS621, demonstrating that the excised intermediate is a covalently closed circular DNA. Left: number of concatemers with the correct LD–RD junction (ACAGTATCTTGTAT; edit distance ≤ 1) across the four conditions (46 vs. 0/0/0). Right: distribution of internal monomer lengths, peaking at the expected excision-product length (~2,579 bp) and distinct from the full-length plasmid (4,638 bp).

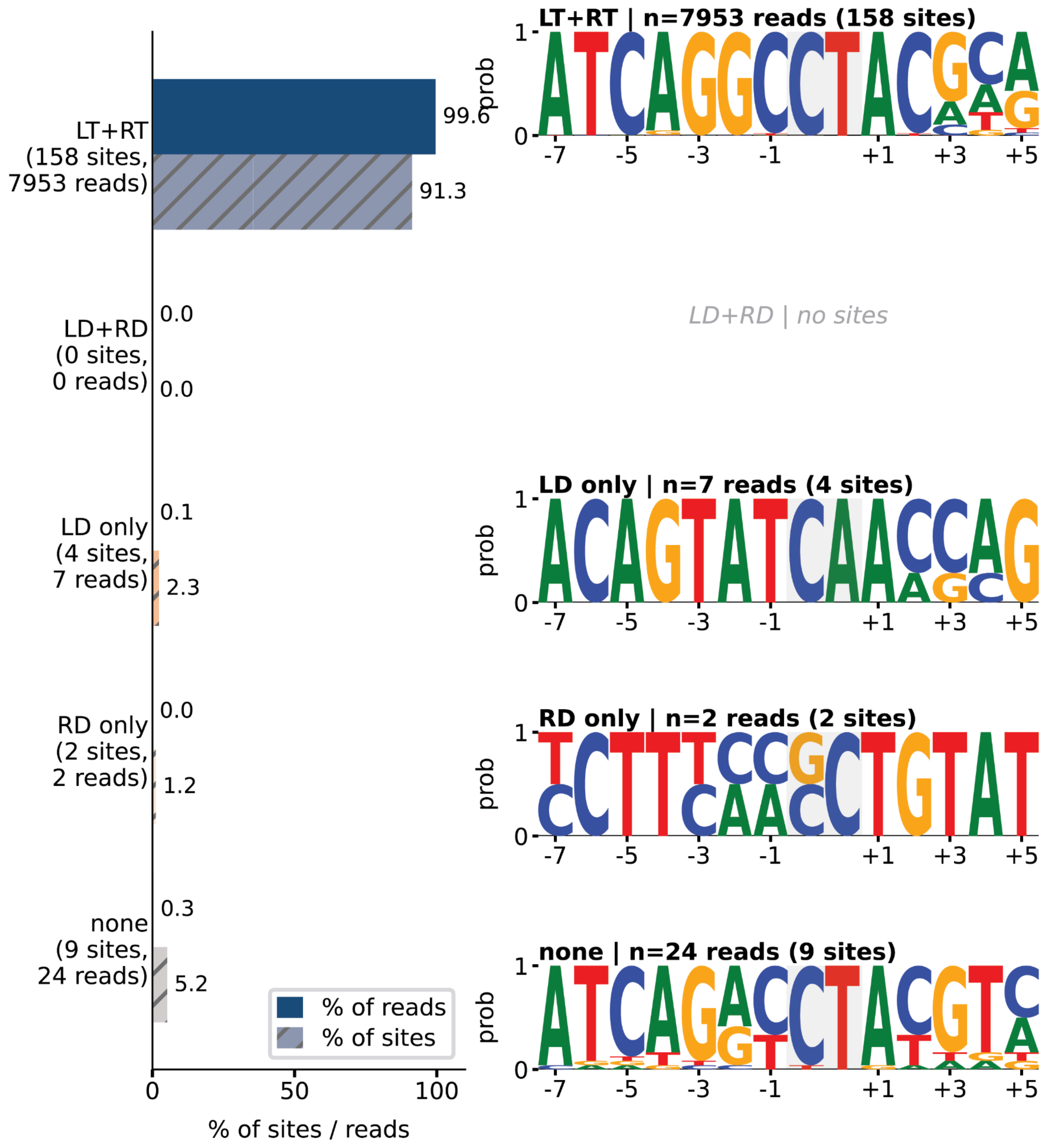


**Fig. S3. Insertion-site recognition categories for the wild-type bridge RNA.**

Insertion sites recovered with the wild type bridge RNA (T-WT/D-WT) were assigned to recognition categories as described in the Methods. Left: percentage of insertion reads (solid) and of insertion sites (hatched) in each category — LT+RT (158 sites, 7,953 reads), LD+RD (no sites), LD-only (4 sites, 7 reads), RD-only (2 sites, 2 reads) and none (9 sites, 24 reads). Right: sequence logos of the 14-bp recombination window recovered in each category. Insertion by the wild-type bridge RNA is almost entirely target-directed: 99.6% of reads and 91.3% of sites fall in the LT+RT category, and donor half-match events (LD-only + RD-only) account for 0.11% of reads and 3.5% of sites.

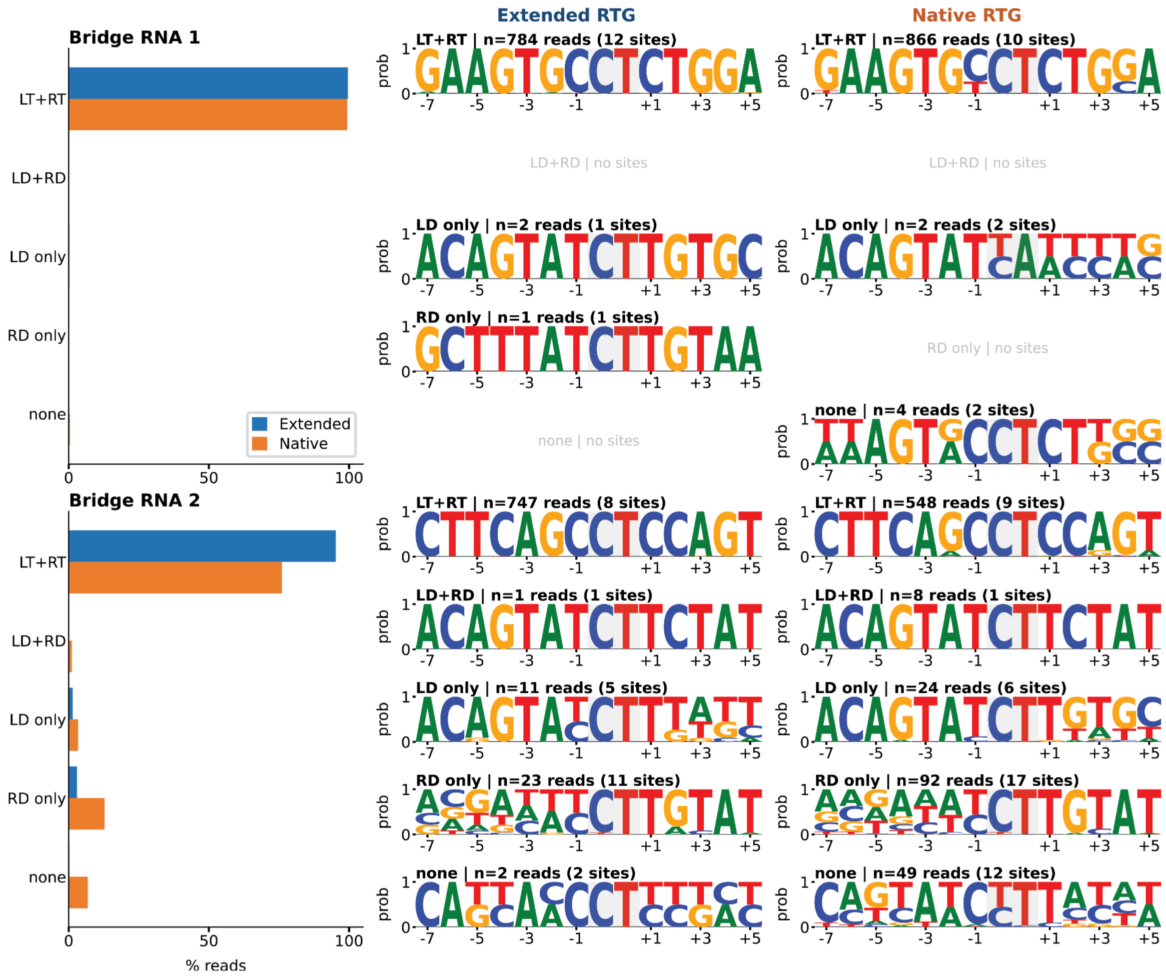


**Fig. S4. Recognition-category distribution for programmed bridge RNAs 1 and 2 with the native and extended RTG.**

Left: percentage of insertion reads in each recognition category for bridge RNA 1 (top) and bridge RNA 2 (bottom), assayed with the native 4-bp right target guide (RTG; orange) and the extended 7-bp RTG (blue). Right: sequence logos of the 14-bp recombination window for each category under each RTG, with the number of reads and sites indicated.

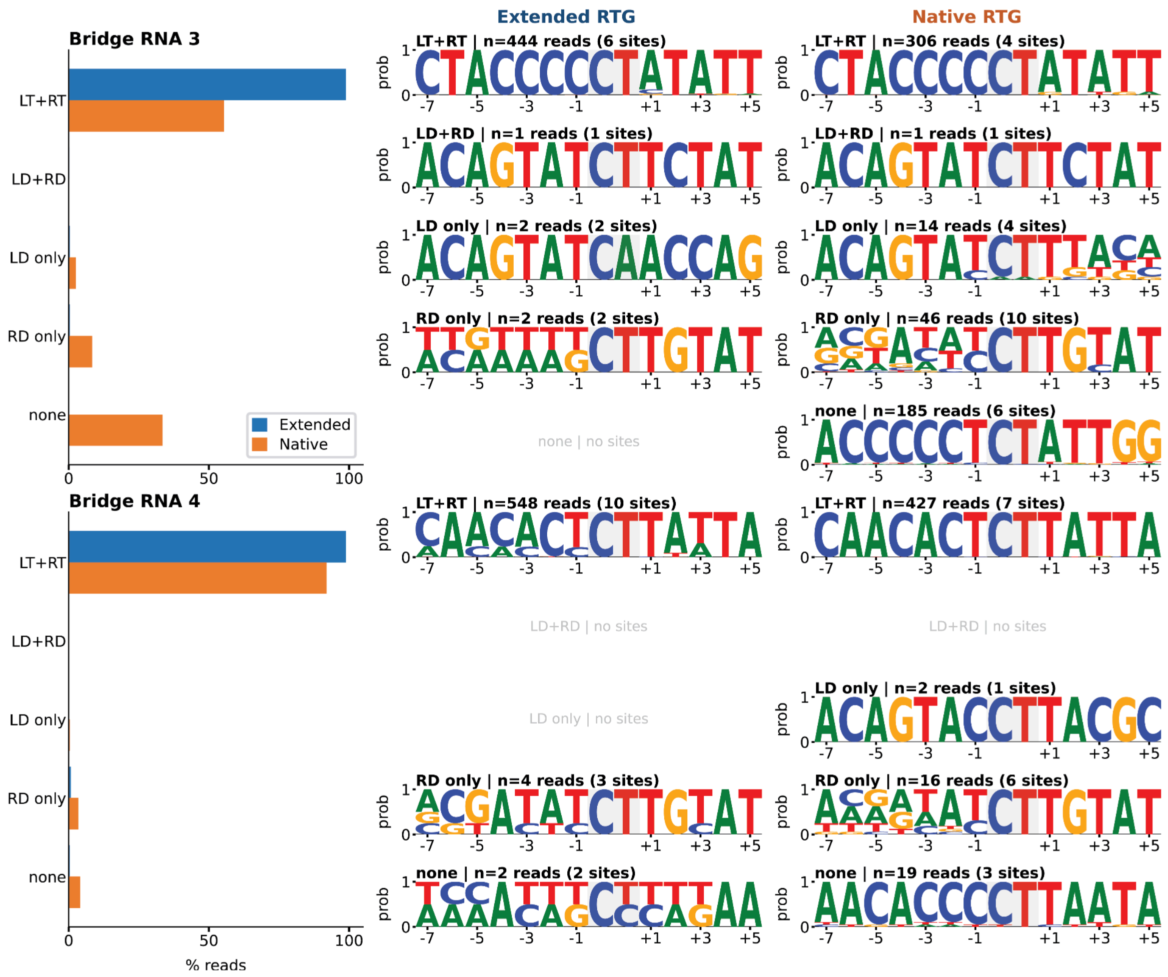


**Fig. S5. Recognition-category distribution for programmed bridge RNAs 3 and 4 with the native and extended RTG.**

Panels as in Fig. S4, for bridge RNA 3 (top) and bridge RNA 4 (bottom).

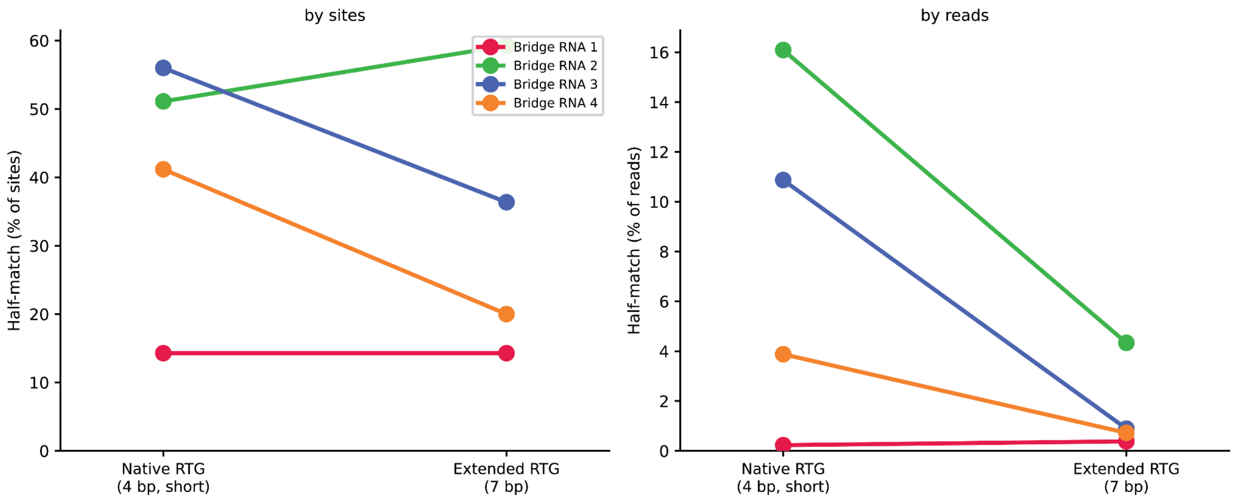


**Fig. S6. Extending the right target guide suppresses half-match insertion.**
Half-match events (LD-only + RD-only) for each programmed bridge RNA with the native 4-bp RTG and the extended 7-bp RTG, expressed as a percentage of insertion sites (left) and of insertion reads (right).

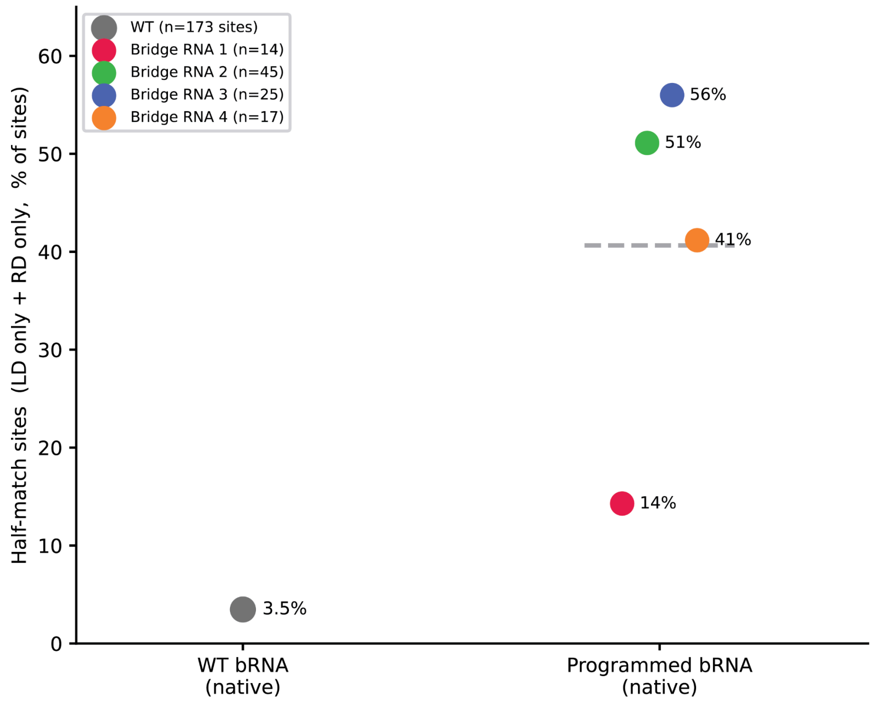


**Fig. S7. Programmed bridge RNAs are more prone to half-match insertion than the wild type bridge RNA.**

Half-match sites (LD-only + RD-only) as a percentage of all insertion sites recovered, comparing the wild type bridge RNA with the four programmed bridge RNAs, all assayed with the native RTG. n, number of insertion sites recovered per guide.
